# An efficient protocol for chromosome isolation from sterlet (*A. ruthenus*) embryos and larvae

**DOI:** 10.1101/2021.05.25.445593

**Authors:** Dorota Fopp-Bayat, Marcin Kuciński

## Abstract

In the present study, the development of an efficient and feasible protocol for chromosome preparation from sterlet (*A. ruthenus*) embryos and larvae was carried out. In the established protocol, the mean efficiency of chromosome extraction ranged from 70 to 100%. The average number of recorded metaphases per slide was between 9 to 15. In general, the most satisfactory results were obtained for embryos at 6 dpf and larvae at the age of up to 7 dph. In the 24 dpf group, chromosome isolation was possible without immersion in spindle poison, however; in successive developmental stages, the minimal immersion time exceeded 1.5 hours, regardless of chorionation. Immersion for 14 hours did not compromise the efficacy of chromosome isolation. In the current study, successful chromosome isolation was determined mainly by hypotonization conditions. Younger developmental stages generally require the shortest hypotonization times, whereas older larvae require longer hypotonization times. The optimal hypotonization period is 5-15 minutes for embryos at 24 dpf, 40 minutes for embryos at 4dpf, and 50-60 minutes for fish at 6 dpf-7 dph. The only exception was the 24 hpf group where only blastula cells were used. An additional overnight fixation step significantly enhanced chromosome quality and supported chromosome counting especially in the 24 dpf group. The quality and quantity of chromosome slides were also significantly determined by tissue type, and the slides prepared from heads (gill cells) produced the best results.

## Introduction

Sturgeons are one of the most valuable fish groups in the global aquaculture, and they are cultured for caviar, tasty meat and isinglass. Sturgeons are also one of the most endangered fish species in the world (FishBase 2021). The development of efficient genome engineering techniques for aquaculture and conservation of sturgeons is a highly promising tool in modern fisheries that continues to attract significant interest (Gui and Zhu 2012; Crego-Prieto et al. 2013; Thresher et al. 2014; Chandra and Fopp-Bayat 2020). Biotechnology tools such as nuclear and mitochondrial DNA markers, gene expression, flow cytometry, genome transplantation and chromosome techniques play a fundamental role in taxonomy, inter-specific hybridization and improvement of commercial fish stocks (Chandra and Fopp-Bayat 2020).

Genome engineering by means of karyotype manipulation encompasses gynogenesis, androgenesis and polyploidization (especially triploidization and tetraploidization). The induction of development in embryos containing uniparental genetic material is known as gynogenesis (maternal material) and androgenesis (paternal material). In aquaculture, both techniques are highly useful for the production of female or male mono-stocks (Chandra and Fopp-Bayat 2020). In sturgeons, gynogenesis plays a particularly important role in caviar production because it reduces the costs associated with the maintenance of spawning stock before it reaches late sexual maturity. Androgenesis is a very useful tool in the restitution and conservation of endangered species in their natural habitat (Corley-Smith and Brandhorst 1999). The triploidization technique plays the key role in polyploidy induction because some triploid fish (especially females) grow at a faster rate and have better tasting meat due to their sterility (Lakra and Das 1998).

Chromosome engineering is also important in developmental biology, reproduction and genetics (Piferrer at al., 2009; Chandra and Fopp-Bayat 2020). It is widely used in research on sturgeons’ evolutionary history, including the determination of differences and abnormalities in ploidy level and the mechanisms responsible for karyotype rearrangement. At present, karyotyping is the only direct method for ploidy determination in sturgeons (Fontana et al. 2003; Fontana et al. 2008; Fopp-Bayat et al., 2017, 2018). Chromosome analysis techniques are also widely applied to study the sex determination system and to identify new sex-related chromosomes in sturgeons (Vasil’eva et al. 2009; Chandra and Fopp-Bayat 2020).

There is considerable evidence to suggest that chromosomes can be effectively isolated from embryos, larvae and adult fish of various species. The isolation of chromosomes from embryos and hatched larvae poses the greatest challenge and requires sensitive procedures (Shao et al., 2010; Ozouf-Costaz et al., 2015; Katami et al., 2015). Despite the availability of various karyotyping techniques such as tissue cultures (Lomax et al., 2000), squashing techniques (Armstrong and Jones 2003) and cell suspensions of mitotic tissues (Henegariu et al., 2001), only one protocol for isolating chromosomes from sturgeons has been developed to date by Fopp-Bayat and Woźnicki (2006), and it relies mainly on the larval stage. Moreover, the extend of optimal chromosome isolation parameters and the impact of fish age on the efficacy of the isolation procedure have never been comprehensively studied. Therefore, the aim of the present study was to develop a feasible, simple and efficient chromosome preparation protocol for sterlet embryos and larvae.

## Results

The developed chromosome preparation protocols enabled the isolation of chromosomes of satisfactory quality, but the results differed across groups. The quality and quantity of chromosome slides were determined by tissue type, and the slides prepared from heads (gill cells) produced the most satisfactory results. The only exception was the 24 hpf group where only blastula cells were used. In the present study, the mean efficiency of chromosome extraction ranged from 70 to 100%. The average number of recorded metaphases per slide was between 9 to 15. The smallest number of chromosomes of the lowest quality were isolated for embryos at the age of 24 hpf, whereas the chromosomes obtained from embryos at 6 dpf and larvae up to 7 dph were of the highest quality (Table 1, Fig. 1). In the current study, the optimal developmental stage for the chromosome preparations was before 7 dph (data not shown).

**Table 1.**
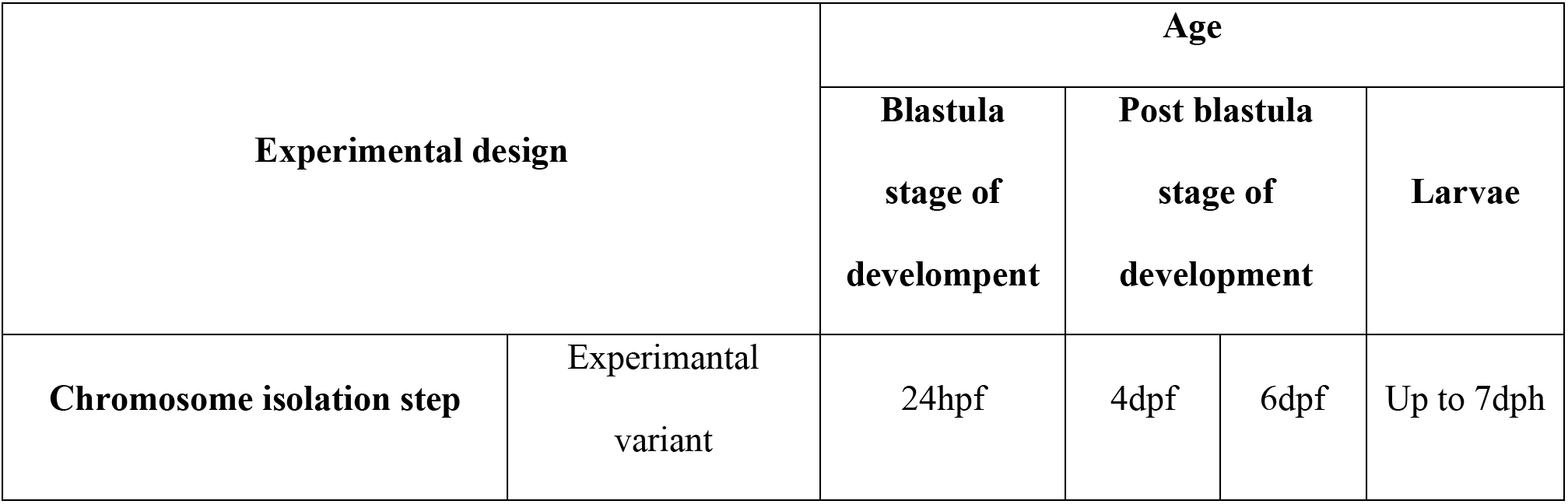
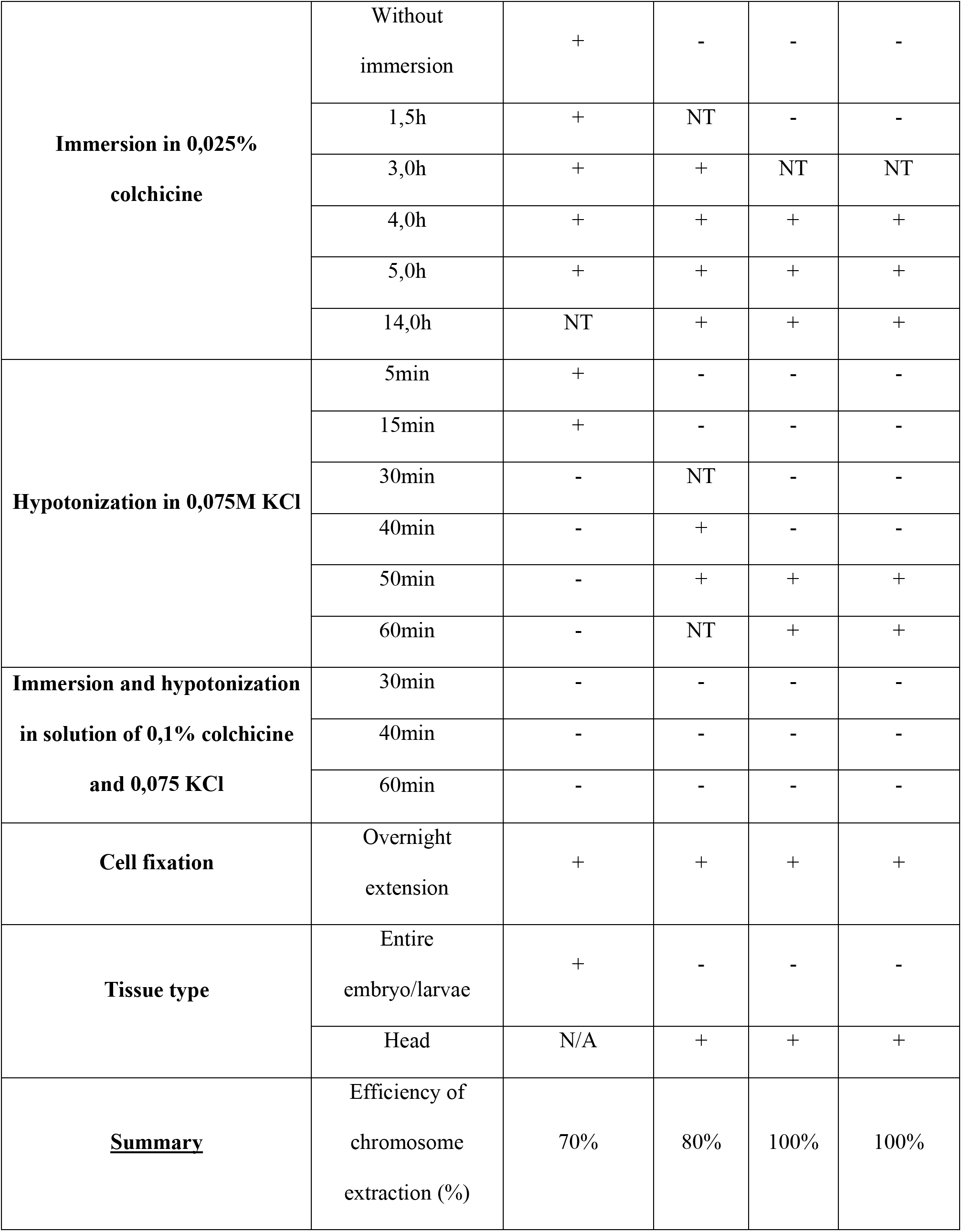
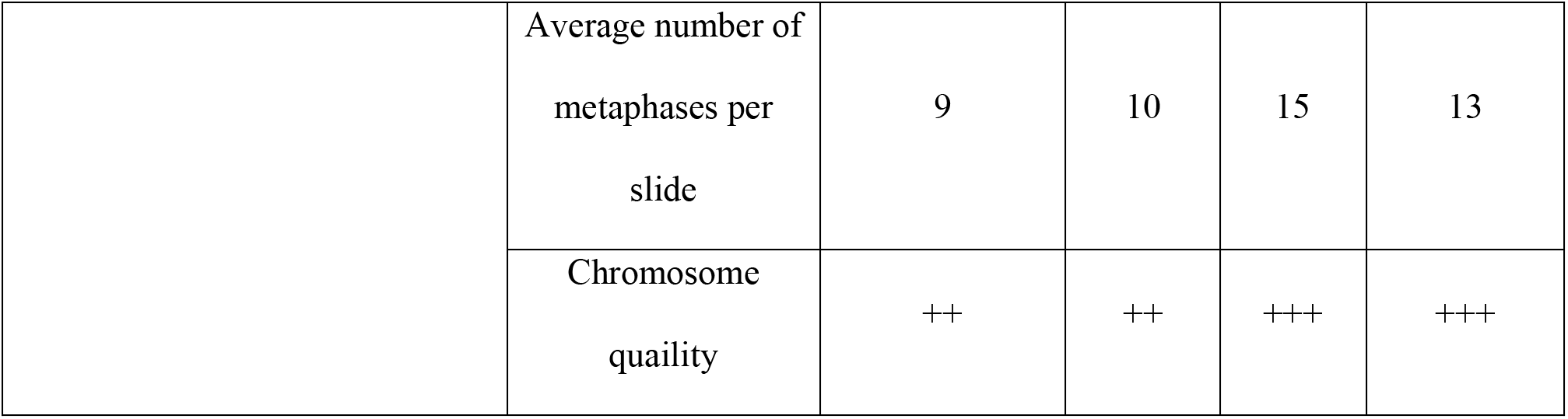
The summary of tested colchicine, hypotonic, fixative treatments, tissue type application and age of fish material utilized together with obtained results. NT: not tested, N/A: not applicable, hpf: hours post fertilization, dpf: days post fertilization, dph: days post hatch. The positive or negative effect of tested variant was marked by plus and minus label, respectively.

| Experimental design |  | Age |  |  |  |
| --- | --- | --- | --- | --- | --- |
|  |  | Blastula stage of development | Post blastula stage of development |  | Larvae |
| Chromosome isolation step | Experimental variant | 24hpf | 4dpf | 6dpf | Up to 7dph |
| **Immersion in 0,025% colchicine** | Without immersion | + | - | - | - |
|  | 1,5h | + | NT | - | - |
|  | 3,0h | + | + | NT | NT |
|  | 4,0h | + | + | + | + |
|  | 5,0h | + | + | + | + |
|  | 14,0h | NT | + | + | + |
| **Hypotonization in 0,075M KCl** | 5min | + | - | - | - |
|  | 15min | + | - | - | - |
|  | 30min | - | NT | - | - |
|  | 40min | - | + | - | - |
|  | 50min | - | + | + | + |
|  | 60min | - | NT | + | + |
| **Immersion and hypotonization in solution of 0,1% colchicine and 0,075 KCl** | 30min | - | - | - | - |
|  | 40min | - | - | - | - |
|  | 60min | - | - | - | - |
| **Cell fixation** | Overnight extension | + | + | + | + |
| **Tissue type** | Entire embryo/larvae | + | - | - | - |
|  | Head | N/A | + | + | + |
| **<u>Summary</u>** | Efficiency of chromosome extraction (%) | 70% | 80% | 100% | 100% |
|  | Average number of metaphases per slide | 9 | 10 | 15 | 13 |
|  | Chromosome quality | ++ | ++ | +++ | +++ |

**Figure 1.**
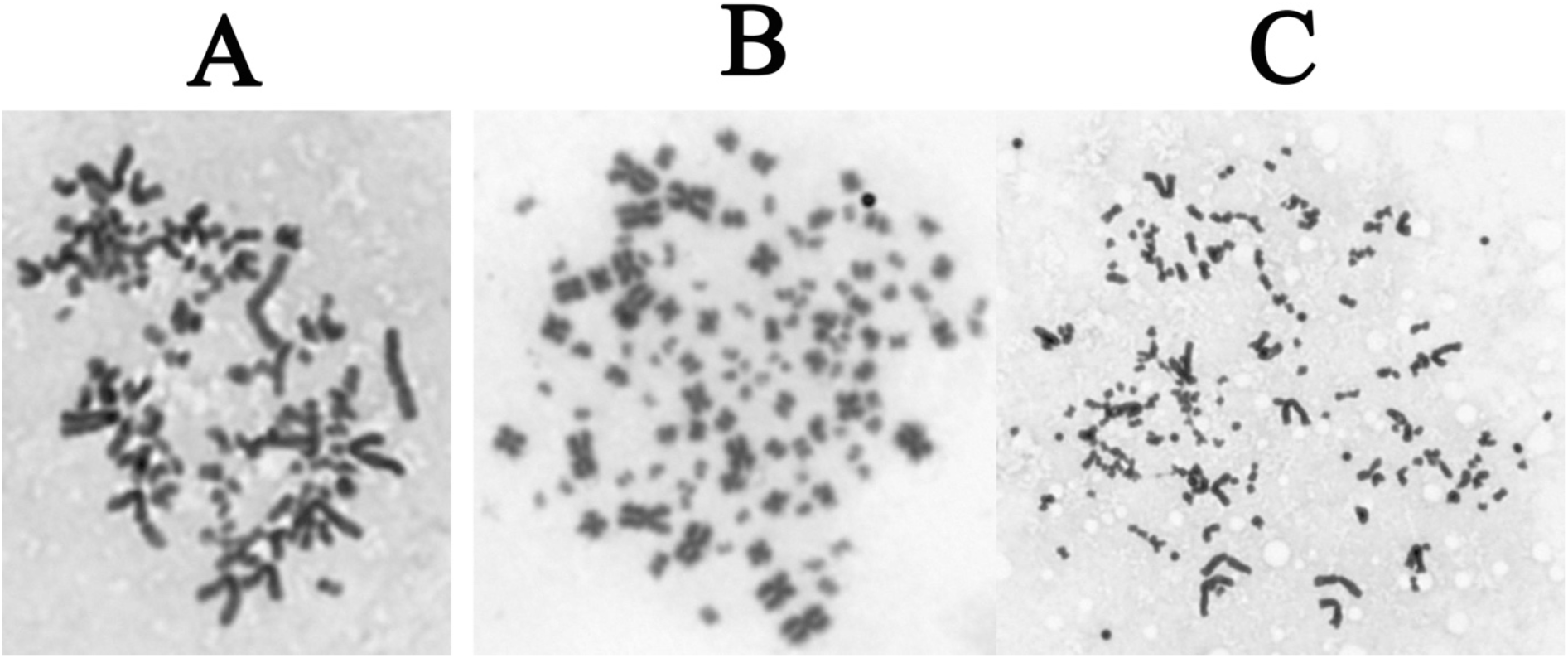
The results of chromosome isolation. The medium and the quality of the isolated chromosomes are marked as A and B, respectively. The result of chromosome isolation without colchicine treatment is marked as C.

In the present study, the incubation time of spindle poison had no significant influence on the effectiveness of chromosome isolation from chorionated or chorion-less embryos. In general, the highest plasticity was observed in embryos at 24 dpf, where positive effects of immersion were noted for every incubation and were associated with the highest rate of mitotic division in this group. In the 24 dpf group, chromosome isolation was possible without immersion in spindle poison that characterized by chromosome slides of relatively high quality. In successive developmental stages, the minimal immersion time exceeded 1.5 hours, regardless of chorionation. Immersion for 14 hours did not compromise the efficacy of chromosome isolation (Table 1).

The optimal hypotonization period was 5-15 minutes for embryos at 24 dpf, 40 minutes for embryos at 4dpf, and 50-60 minutes for fish at 6 dpf-7 dph. The alternative step combining immersion and hypotonization failed to generate positive results (Table 1).

The applied 12-14h additional fixation step significantly improved the quality of the isolated chromosomes. The most pronounced improvement was observed in the 24 hpf group, which significantly enhanced chromosome quality and supported chromosome counting. In turns, the least noticeable improvement was recorded in 6 dpf-7 dph groups.

## Discussion

The majority of chromosomal isolation protocols have been designed for specific species, and the effects of each step in the procedure of isolating chromosomes from adult fish or larvae fish, in particular the omission of embryonic developmental stages, have been examined by different authors. In the present study, an optimal protocol was developed for isolating chromosomes from the embryos and larval developmental stages of *A. ruthenus*. The results supported the identification of three main chromosome preparation approaches that differ in the parameters of each step for fish at: (1) 24 hpf, (2) 4 dpf and (3) 6 dpf-7 dph (Table 1; Fig. 1 and 3).

**Figure 2.**
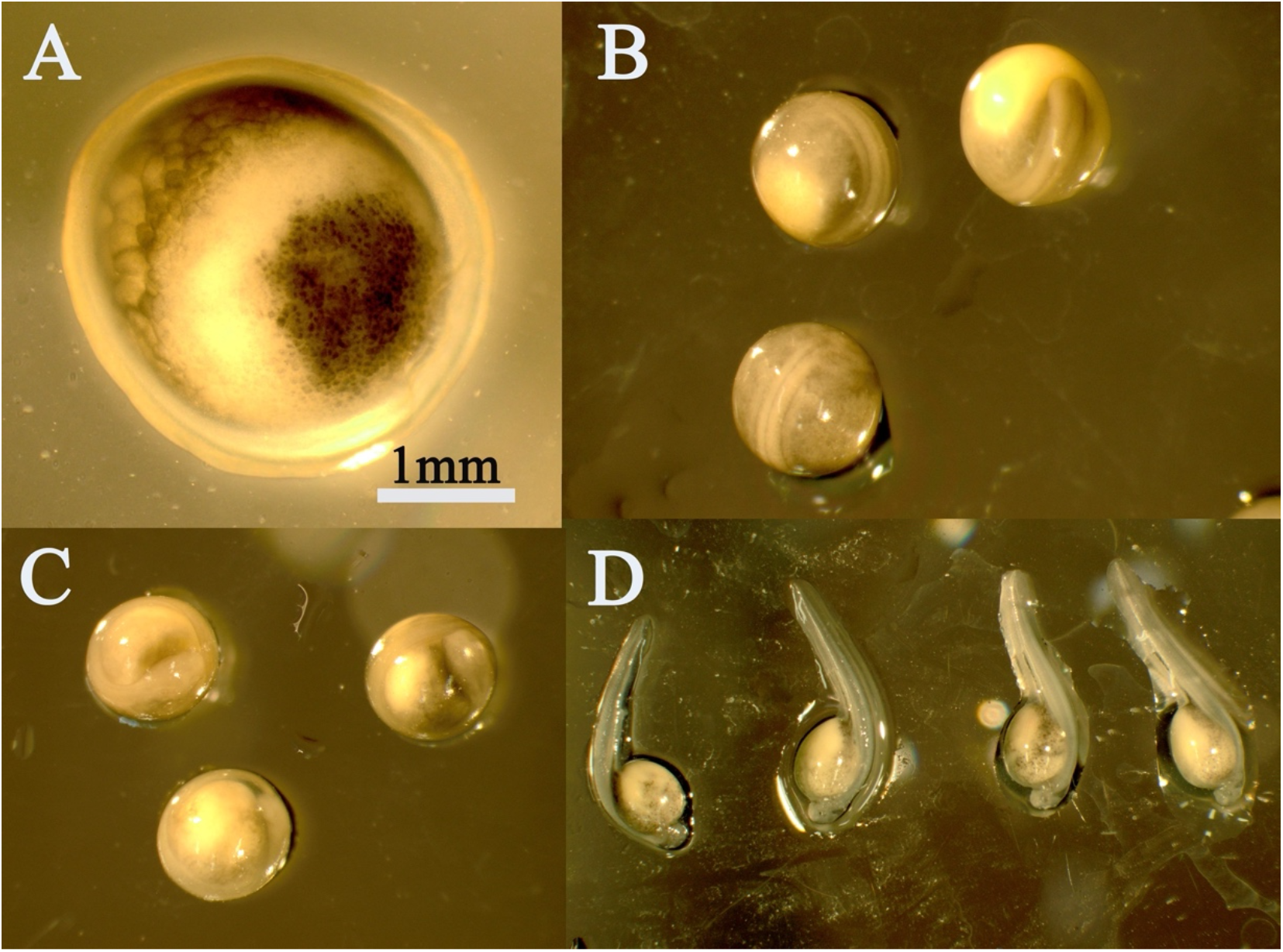
Fish material used in the study. A: embryos at 24 hours post-fertilization (hpf) (blastula stage), B: 4 days post-fertilization (dpf), C: 6 dpf (before hatching), and D: larvae with the yolk sac up to 7 days post-hatching (dph).

**Figure 3.**
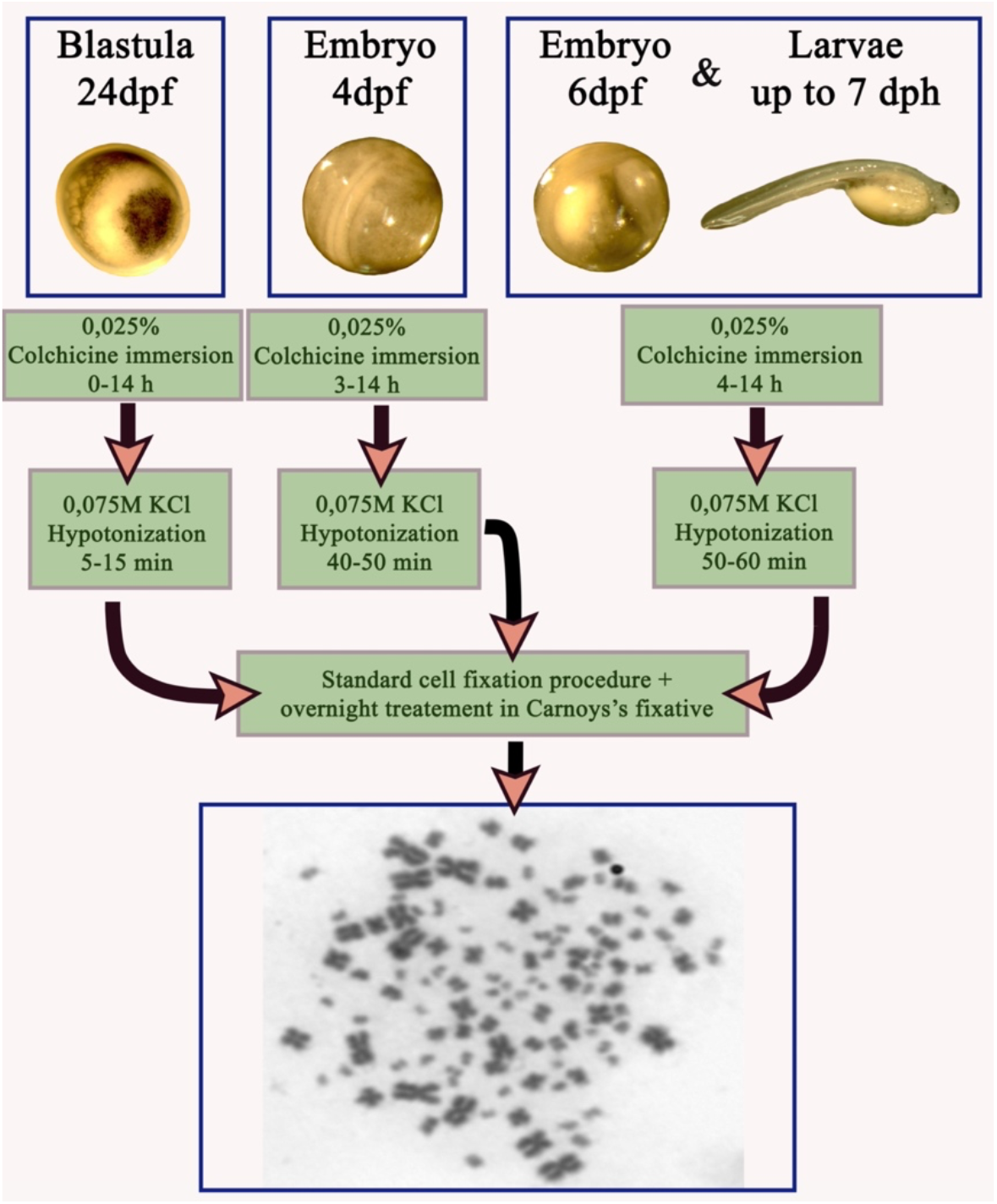
The graphical summary of determined three main chromosome isolation protocols for sterlet embryos and larvae.

In most studies, the achievement of clear and identifiable metaphase chromosome spreads requires sufficient concentrations and incubation times of tissues in the spindle poison (Rieder and Palazzo 1992). In general, insufficient concentration and/or duration of exposure to the spindle poison prevents the interception of metaphase cells, whereas very high concentrations and/or prolonged exposure may lead to chromosomal condensation (Wood et al., 2001). The most common microtubular poison for cell interception is colchicine which inhibits spindle microtubules before envelope breakdown (NEB) in metaphase cells (Karami et al., 2015). In other studies, fish age was also correlated with minimal immersion time, which implies that the procedure should be optimized for individual species and age groups (Caperta et al., 2006; Karami et al., 2015). However, according to some authors, killing the larvae and separating the yolk before incubation in colchicine is a more efficient procedure than incubating live larvae (Hussain and McAndrew 1994; Pradeep et al., 2011; Karami et al., 2015). These authors also hypothesized that yolk size and absorption rate are the key facts that differentiate chromosome preparation protocols.

After mitotic spindle inhibition, cells and tissues must be incubated in a hypotonic solution to swell the nuclei and spread the chromosomes on the slides (Moore and Best 2001). The hypotonic solution and incubation time have to be optimized to prevent chromosome knotting, overlapping or its partial loss (Baksi and Means 1988). According to some authors, the yolk should be separated from hypostasized tissues because its high lipophilicity can prevent the hypotonic solution from penetrating the cells (Baksi and Means 1988). Potassium chloride (KCl 0.075 M) is one of the most widely used hypotonic solutions. In the current study, successful chromosome isolation was determined mainly by hypotonization conditions. Younger developmental stages generally require the shortest hypotonization times, whereas older larvae require longer hypotonization times (Table 1). Other authors demonstrated that distilled water can produce superior results, which suggests that the type of hypotonic solution should be carefully selected for the analyzed species and age group (Karami et al., 2015).

In the next obligatorily step of the chromosome preparation protocol, samples are fixed in Carnoy’s fixative solution and a cell suspension is prepared. In this step, the evaporation of the fixative solution and slide preheating can affect the quality and quantity of the isolated chromosomes (Moore and Best 2001). In general, this step is relatively similar in all chromosome isolation protocols for fish. Due to the high effectiveness of this procedure, additional modifications were not required, and the only variation was a 12-14 fixation step which significantly improved the quality of the isolated chromosomes. However, the observed improvement varied across groups and was most pronounced in the 24 hpf group and least noticeable in 6 dpf-7 dph groups. In the 24 dpf group, an additional overnight fixation step significantly enhanced chromosome quality and supported chromosome counting.

Fish transport and other handling operations exerted a negative effect on chromosome preparation efficacy. Embryos and larvae should not be acclimatized after transport or other significant manipulations because the number of the isolated chromosomes declines steadily over time. The above can be attributed to stress that inhibits cell proliferation (Mohanty et al., 2018). Therefore, chromosome preparations should be made immediately after fish transport and handling. It should also be noted that larval age is yet another critical determinant of chromosome isolation efficacy. The effectiveness of chromosome isolation generally decreases on successive days after hatching because the mitotic rate is affected by species, age and environmental conditions (Wakahara 1972; Shelton et al., 1997; Shao et al., 2010). In African sharptooth catfish (*Clarias gariepinus*) and zebra fish (*Danio rerio*), the samples isolated at 4 dph failed to produce countable mitotic chromosome spreads (Karami et al., 2015). However, this threshold time is species-specific and needs to be experimentally determined. In the current study, the optimal developmental stage for the chromosome preparations was before 7 dph (data not shown).

In summary, this study demonstrated that chromosomes can be isolated in every developmental stage of *A. ruthenus*, but the overall quality and quantity of the extracted chromosomes can vary. Under the current study three main chromosome isolation procedures were determined (Fig. 3). In general, the most satisfactory results were obtained for embryos at 6 dpf and larvae at the age of up to 7 dph. In both groups, the quality of the isolated chromosomes supported counting and karyotyping. The chromosomes isolated from 24 hpf and 4 dpf groups were characterized by the lowest quantity and quality. These isolates could be used for chromosome counting only, and high-quality metaphases for karyotyping were rarely obtained.

## Materials and Methods

A series of experiments based on the chromosome preparation procedure described by Fopp-Bayat and Woźnicki (2006), which is a modified version of the spindle poison technique developed by Kligerman and Bloom (1977), were conceived and conducted to establish the chromosome isolation protocol. To this end, ten embryos and larvae of *A. ruthenus* were used in each experimental variant. The fish material differed in age to determine the influence of developmental stage on the effectiveness of chromosome isolation. Four age variants were used: (1) embryos 24 hours post-fertilization (hpf) (blastula stage), (2) 4 days post-fertilization (dpf), (3) 6 dpf (before hatching), and (4) larvae with a yolk sac up to 7 days post-hatching (dph) (Table 1 and Fig. 1). Fish material was obtained from a fish farm in Wąsosze (southern Poland), and the study was performed in the Center of Aquaculture and Environmental Engineering in Olsztyn (north-east Poland). The experiments were performed immediately after the delivery of the fish material.

The experimental chromosome preparation procedure comprised five main steps, where all variables were optimized in a stepwise manner (Table 1). The following variants were tested:

1. Immersion of live embryos (chorionated or chorion-less) and larvae in 0.025% colchicine at 16°C for 1.5, 3.0, 4.0, 5.0 and 14.0 hours (h). The variant was also tested without immersion,
2. Hypotonization of embryos and larvae in 0.075 M solution of KCl for 5, 15, 30, 40, 50 and 60 minutes (min),
3. Immersion and hypotonization in a solution of 0.1% colchicine and 0.075 KCl for 30; 40 and 60 minutes was tested simultaneously as an alternative to the first and second step,
4. Tissue fixation in cooled Carnoy’s fixative solution (methanol and acetic acid, 3:1). A fresh solution was prepared each time, and it was triple fixed at −20°C in every experimental variant. Each fixation stage lasted 30, 15 and 15 minutes. A fourth fixation step of 12-16 hours (overnight) was additionally applied to check the effect of extended fixation on the quality of the isolated chromosomes,
5. Homogenization of different types of tissues. Blastula cells extracted from the chorion and separated from the yolk sac were used in the homogenization of fish material at 24 hpf. Tissues from the entire embryo/larva and head tissues were used in the remaining age groups (Table 1). Tissues were homogenized with a needle in Eppendorf tubes containing a small volume (around 0.1-0.2 ml) of fresh fixative solution.

The tissues intended for chromosome preparations were carefully separated from the yolk sac after colchicine immersion. Chromosome slides were prepared in accordance with the standard splash procedure, where several droplets (1-3) of the homogenized cell suspension are dropped onto microscope glass slides from a height of around 30 cm. Before chromosome preparation, the glass slides were washed in solution of 96% ethanol containing several drops of condensed hydrochloric acid (HCl) (Fopp-Bayat and Woźnicki 2006). One or two slides were prepared from each embryo or larva.

The quality of the obtained chromosome slides was checked under the Zeiss Axio Imager.A1 microscope equipped with a digital camera. Chromosome slides were examined under bright light, and the quality of the extracted chromosome was classified into three main categories: (1) high quality results characterized by a large number of slides with successfully isolated chromosomes (above 80%) that are eligible for counting and karyotyping (+++), (2) medium quality results characterized by an average number of slides with successfully isolated chromosomes (30-80%) that can be used only for counting, and (3) no results or low quality results characterized by a small number of slides containing chromosomes (below 30%) that cannot be used for cytogenetic analysis. Clear chromosome spreads were counted on each slide to assess the quantity of the chromosomes isolated in each experimental variant.

## Acknowledgements

We would like to thank Elzbieta and Andrzej Fopp from Wasosze Fish Farm for providing the sterlet embryos for experiments. We also thank Gyan Chandra and Paweł Woznicki for laboratory assistance and valuable suggestions. We would like to also thank to anonymous reviewers whose valuable suggestions essentially contributed to improvement of the present manuscript.

